# On-farm Implementation of Midseason Drainage to Decrease Greenhouse Gas Emissions and Grain Arsenic Concentration in Rice Systems

**DOI:** 10.1101/2022.03.23.485547

**Authors:** Henry Perry, Daniela R. Carrijo, Aria H. Duncan, Scott Fendorf, Bruce Linquist

**Author notes:** Corresponding author: Henry Perry, University of California, 1 Shields Ave, Davis, CA, USA.

## Abstract

Rice (*Oryza sativa L.*) is an important staple crop throughout much of the world, however, it is also a significant source of agricultural methane (CH_4_) emissions and exposure to arsenic (As). Introduction of soil aerobic events through practices such as alternate wetting and drying or midseason drainage, in flooded rice systems can significantly decrease grain As concentration and seasonal CH_4_ emissions. Previous on small research plots research has shown that a single midseason drain accomplishes these goals without yield reduction, but the degree of benefit depends on soil-drying severity. A midseason drain also has the potential to fit in well within current management practices of California rice systems, however, it has not been tested across a wide range of soil types or at a scale that farmers typically manage in this region. Therefore, in this three-year study, we aimed to determine if the results from previous small plot research are similar to what can be expected on-farm. At seven on-farm trials we implemented a single midseason drain and compared the grain yields, GHG emissions, and As concentration to the traditional farmer practice (FP) practiced in an adjacent part of the field. Soil moisture parameters [perched water table, volumetric water content, gravimetric water content (GWC), and soil water potential], CH_4_ and nitrous oxide (N_2_O) emissions, grain As and cadmium concentration, and grain yield were quantified. Across site-years, midseason drainage reduced seasonal CH_4_ emissions by 20-77%, compared to the FP control with the magnitude of reduction related to the soil-drying severity. For every 1% reduction in soil GWC during the drainage period, CH_4_ emissions were reduced by approximately 3.2%, compared to 2.5% in previous on-station research using small plots. Midseason drainage increased N_2_O emissions (average = 0.248 kg N_2_O-N ha^-1^) compared to the control but this accounted for only 3% of the seasonal global warming potential across all drainage treatments. Drainage also decreased grain As concentration by approximately 20%, on average, but was not related to the degree of soil-drying. Importantly, midseason drainage had no significant impact on grain yields. Overall, these results confirm findings from previous on-station research, indicating that midseason drainage may be a viable on-farm management practice for GHG mitigation and for reducing grain As concentration in flooded rice fields with limited risk of yield reduction.

## 1. Introduction

Rice is a major staple crop for nearly half of the world’s population, however, it is also a significant source of agricultural greenhouse gas (GHG) emissions, particularly methane (CH_4_). In fact, rice cultivation is responsible for more than 20% of agriculturally related CH_4_ emissions (IPCC, 2013), and the global warming potential (GWP) of rice is approximately 2.5-5.5 times higher than that of the other major cereal crops (Linquist et al., 2012). Additionally, rice can be a significant source of exposure to arsenic (As), which is toxic to humans, especially in its inorganic form, and is of particular concern in countries in which rice consumption is high (Meharg, 2004; Bhowmick et al., 2018).

In most rice systems, fields are kept flooded for the majority of the growing season, which creates anerobic conditions in the soil. Such conditions are favorable for elevated CH_4_ emissions as well as increased soil As mobilization and plant uptake. Methane in particular is the product of decomposition of organic matter under anaerobic soil conditions (Conrad, 2007). Many studies have explored the effects of alternative water management practices on GHG emissions in flooded rice systems, and such practices are commonly known by several different names including alternate wetting and drying (AWD), intermittent irrigation, or midseason drainage. Midseason drainage refers to the management practice in which a single dry-down event is employed during the mid-vegetative to early reproductive stages of rice growth. This is in contrast with AWD or intermittent irrigation practices, in which two or more drainage events are generally employed throughout the season. While midseason drainage and AWD have been shown to significantly decrease CH_4_ emissions, N_2_O emissions can often increase as a result of these practices (LaHue et al., 2016; Perry et al., 2022; Zou et al., 2007) and are the result of soil nitrification and denitrification processes during soil-drying and reflooding, respectively (Bateman and Baggs, 2005).

In addition to GHG mitigation benefits, alternative water management practices such as midseason drainage have also been utilized to effectively decrease grain As concentration in flooded rice systems (Lahue et al., 2016; Linquist et al., 2015; Carrijo et al., 2019). Anaerobic soil conditions lead to the reduction of As(V) (arsenate) to more mobile As(III) (arsenite) and the reductive dissolution of As-bearing iron (Fe) (hydr)oxides (Fendorf et al., 2007). Soil-drying sequester As. However, cadmium (Cd) bioavailability tends to increase under soil aerobic conditions, primarily due to desorption with decreasing soil pH, resulting in uptake by rice when drainage occurs (Rinklebe et al., 2016; Wang et al., 2019). Studies evaluating the effect of non-continuous flooding on grain As and Cd are lacking on-farm. Therefore, it is critically important to scale up these practices to commercial rice fields to determine if the results generated from small-plot experiments can be expected at field-scale.

In a review by Jiang et al. (2019), the authors reported that, on average, non-continuous flooding practices decreases seasonal CH_4_ emissions by approximately 53%, with single and multiple drainage events reducing CH_4_ emissions by an average of 33% and 47%, respectively. In general, increased soil-drying severity during imposed drainage periods further reduces CH_4_ emissions (Balaine et al., 2019; Jiang et al., 2019), however, this exact relationship is relatively unknown across a variety of soil types and in different environmental conditions. Additionally, the majority of studies which have implemented non-continuous flooding in rice systems have been conducted in small, controlled plots (de Vries et al., 2010; Dunn and Gaydon, 2011) or at field stations (LaHue et al., 2016; Balaine et al., 2019), in which the size of experimental units is often significantly less than that of traditional commercial rice fields, and unforeseen complications are usually minimized. Importantly, managing irrigation of rice at a large scale can be challenging and may significantly affect GHG emission estimates due to heterogeneity of soil moisture when draining and reflooding fields.

With regards to the effects of midseason drainage and other non-continuous flooding practices on grain yield, many studies have found that these practices have no significant impact on yield (Balaine et al., 2019; Lu et al., 2000; Pandey et al., 2014; Perry et al., 2022; Yao et al., 2012). In general, the number of drainage events has also been found to not be a significant factor in determining grain yield, however, increased soil-drying severity can significantly reduce yields (Jiang et al., 2019). Overall, non-continuous flooding practices have been found to decrease grain yield by 3.6%, on average (Jiang et al., 2019), however, the precise yield-response to drainage depends heavily on a variety of factors such as soil type, rice cultivar, soil-drying severity of the drainage period(s), and specific environmental conditions and characteristics. Importantly, in locations in which the water table is relatively shallow as was the case in Balaine et al. (2019) and Perry et al. (2022), rice plants can tolerate relatively high levels of soil-drying as roots can grow deeper into the soil profile to reach the shallow water table, aiding in water uptake during drying periods (Carrijo et al., 2018). Conversely, in locations in which the water table is relatively deep, rice may not be able to tolerate high levels of soil-drying. Additionally, periods of non-continuous flooding can often coincide with the occurrence of rice blast, a harmful fungal pathogen caused by *Magnaportha grisea*, for cultivars which are not blast-resistant, resulting in significant yield losses (Nalley et al., 2014). That said, modern hybrid cultivars are blast-resistant and capable of withstanding periods of moderate to severe soil-drying, therefore non-continuous flooding practices such as midseason drainage should ideally be implemented using modern blast-resistant rice cultivars.

Importantly, while soil-drying events can decrease plant uptake of As, Cd uptake can increase under soil aerobic conditions, and several studies have shown that non-continuous flooding has both decreased total grain As concentration and increased grain Cd concentration (Li et al., 2019; Martinez-Eixarch et al., 2021; Norton et al., 2017). Decreases in total grain As concentration across studies are also quite variable, although increased soil-drying severity generally leads to further decreases in total grain As concentration.

There are a number of challenges associated with implementing non-continuous flooding practices on-farm. During the wet season in Asian rice systems, for example, midseason drainage may not be feasible as fields may not drain properly and can lead to limited GHG mitigation potential compared to drainage in the dry season (Sander et al., 2017). Additionally, many surface irrigation schemes in Asia consist of decades-old canal infrastructure which lead to substantial water losses and can limit water availability when it is needed for reflooding (Barker and Molle, 2004). Groundwater pumping, which is less common in these areas, can mitigate water supply issues but requires substantial energy input and can also lead to ecological degradation (Khan et al., 2006). Furthermore, groundwater in many rice producing regions is highly contaminated with As, and successive cycles of irrigation with As-laden groundwater can lead to further accumulation of As in paddy soils (Dittmar et al., 2007).

In California most rice is irrigated using surface water and is delivered through irrigation districts, which can limit growers’ flexibility of applying water when it is needed. In California rice systems, water management is also primarily driven by herbicide applications and water holding time requirements after herbicide applications, limiting the flexibility of when water can be drained from the field. In addition to the variability of water dynamics for large-scale flooded rice fields, slow adoption of alternative water management amongst farmers due to traditional beliefs and habits and the aforementioned limitations associated with flexibility in water management are significant obstacles for wide-spread adoption of non-continuous flooding

However, there are a few reasons why midseason drainage in particular may be a practical on-farm alternative to practices which employ multiple drainage periods. Firstly, a single dry-down event requires less overall maintenance and less risk of yield loss, on average, than practices which employ several drainage events throughout the season (Carrijo et al. 2017; Jiang et al., 2019). As mentioned earlier, in California rice systems it can also be particularly difficult to manage multiple drainage events on-farm due to strict water-holding time requirements following pesticide applications. In the Sacramento Valley of California, in which 95% of the state’s rice is grown (UCANR 2021), it is also common for rice growers to apply a midseason application of the contact herbicide, propanil, for weed suppression, the process of which involves draining floodwater to expose aquatic weeds prior to application and subsequent reflooding shortly after application. For these reasons, midseason drainage for GHG mitigation may align well with many CA rice growers’ current management practices and could be well-suited for California rice cultivation as a whole.

In California as well as in the mid-southern United States, rice farms are, on average, larger than in Asia, the rice bowl of the world. In California, the average rice farm is approximately 200 ha (Childs et al., 2020), while in Asia the size of the average rice farm ranges between 0.5-2.0 ha (USDA 2015). Rice fields are also generally smaller in Asia than in California. Importantly, and with regards to water management, in the United States, rice fields are often divided into multiple basins or checks, in which water is supplied serially from the topmost to the bottommost check and is regulated by adjustable weirs or rice boxes (Hill et al., 1991). In this three-year study, we aimed to scale up midseason drainage to individual checks of conventionally managed rice fields in the Sacramento Valley of California.

In a previous on-station study using small plots (0.3 ha) we found that a single midseason drainage period of varying soil-drying severity and timing decreased seasonal CH_4_ emissions between 38-66% without significantly impacting yields (Perry et al., 2022). In this study, our broad objective was to implement a midseason drain on-farm across a wide range of grower management practices and soil types to determine if the results we found in small on-station plots are similar to what can be expected in commercial California rice fields. Our specific objectives were to assess the variability of check-scale soil-drying and to quantify across a variety of soil types and management practices the effect of soil-drying severity on grain yield, grain As concentration, and CH_4_ emissions. We hypothesized that the relationship between soil-drying severity during the drainage period and percent reduction in cumulative seasonal CH_4_ emissions compared to the control would be similar to that which we had determined from our previous on-station study. We also hypothesized that midseason drainage would not significantly affect grain yield compared to the control and that drainage would significantly decrease total grain As concentration while increasing grain Cd concentration, compared to the control.

## 2. Materials and Methods

### 2.1 Study site description

Seven on-farm trials were established during the 2017, 2019, and 2020 rice growing seasons at various locations (referred to by proximity to the nearest town and the study year) throughout the Sacramento Valley in California. This region has a Mediterranean climate, characterized by warm and dry conditions during the rice growing season from May to October. The average air temperature and average total precipitation across the three growing seasons in this study period were 22.9 °C and 40.4 mm (CIMIS Biggs 2021), and there was no precipitation during the drainage periods. Soil samples were collected from the plow layer (approximately 0-15 cm) after tillage and before fertilizer application. Soil taxonomic classification as well as selected chemical and physical properties for each site-year are provided in Table 1. Most of the study-sites consisted of soils with relatively high clay content (38-55%), which is typical of rice soils in California. The two exceptions to this were Marysville-19 (25%) and Marysville-20 (22%). Soil pH, measured using saturated paste (United States Salinity Laboratory Staff, 1954), ranged from 4.6 to 7.0. Soil organic C and N, measured using an elemental analyzer interfaced to a continuous flow isotope ratio mass spectrometer (EA-IRMS), ranged from 1.33 to 2.34% and 0.11 to 0.20%, respectively.

**Table 1.** Soil description, characteristics, and selected properties of each site-year.

| Site-year | Soil series | Taxonomic classification | Total Arsenic | Total Cadmium | Texture |  |  | Organic carbon | Total nitrogen | pH |
| --- | --- | --- | --- | --- | --- | --- | --- | --- | --- | --- |
|  |  |  | ppm | ppm | Sand | Silt | Clay | % | % |  |
| Elverta-17 | Clear Lake Clay | Fine, smectitic, thermic Xeric Endoaquerts | 0.34 | 6.5 | 27 | 33 | 40 | 2.34 | n/a | 5.3 |
| Marysville-19 | San Joaquin Loam | Fine, mixed, active, thermic Abruptic Durixeralfs | 6.62 | 2 | 32 | 43 | 25 | 1.56 | n/a | 5.1 |
| Willows-19 | Castro Clay | Fine, thermic Typic Calciaquolls | 6.26 | 2 | 26 | 36 | 38 | 2.06 | n/a | 7.0 |
| Arbuckle-20 | Clear Lake Clay | Fine, smectitic, thermic Xeric Endoaquerts | 9.69 | 3 | 10 | 35 | 55 | 2.24 | 0.20 | 6.4 |
| Willows-20 | Castro Clay | Fine, thermic Typic Calciaquolls | 6.07 | 2 | 28 | 34 | 38 | 2.09 | 0.18 | 6.8 |
| Marysville-20 | San Joaquin Loam | Fine, mixed, active, thermic Abruptic Durixeralfs | 2.84 | 1.5 | 38 | 40 | 22 | 1.34 | 0.11 | 4.6 |
| Pleasant Grove-20 | Capay silty clay | Fine, smectitic, thermic Typic Haploxererts | 8.32 | 2 | 17 | 34 | 49 | 1.33 | 0.13 | 6.6 |

### 2.2 Field management

All fields received applications of pre-plant mineral N fertilizer in the form of aqua ammonia at varying rates and were aerially water-seeded after seedbed preparation, fertilization, and flooding (Table 2). After N fertilization and flooding, fields were direct seeded with either rice variety M-105, M-206, or M-209. Herbicide and pesticide applications occurred during the first 6 weeks of crop development as is common practice in California rice systems. About 3 weeks before harvest all plots were drained and allowed to dry in preparation for harvest. At the five site-years in which GHG emissions were measured, rice straw from the previous season’s harvest was either burned, chopped and incorporated into the soil followed immediately by winter-flooding, or the field was fallowed in the previous growing season.

**Table 2.**
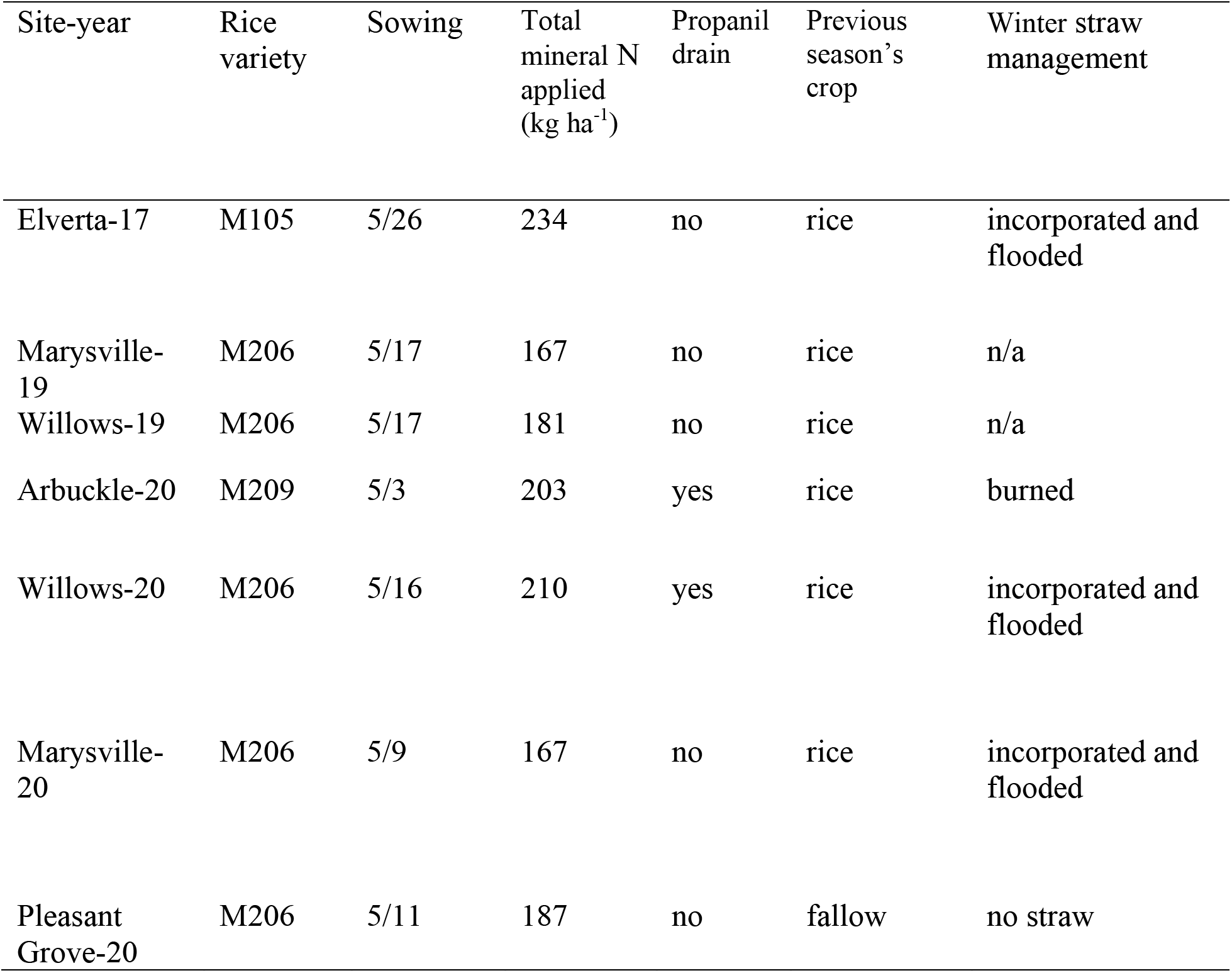
Field and crop management information.

| Site-year | Rice variety | Sowing | Total mineral N applied (kg ha<sup>-1</sup>) | Propanil drain | Previous season's crop | Winter straw management |
| --- | --- | --- | --- | --- | --- | --- |
| Elverta-17 | M105 | 5/26 | 234 | no | rice | incorporated and flooded |
| Marysville-19 | M206 | 5/17 | 167 | no | rice | n/a |
| Willows-19 | M206 | 5/17 | 181 | no | rice | n/a |
| Arbuckle-20 | M209 | 5/3 | 203 | yes | rice | burned |
| Willows-20 | M206 | 5/16 | 210 | yes | rice | incorporated and flooded |
| Marysville-20 | M206 | 5/9 | 167 | no | rice | incorporated and flooded |
| Pleasant Grove-20 | M206 | 5/11 | 187 | no | fallow | no straw |

### 2.3 Treatments and experimental design

In each year of the study, two water management treatments—midseason drainage (MD) and a traditional farmer practice (FP) control—were tested in individual rice field checks, ranging between 2-8 ha. The checks of rice fields were separated by levees, and water was supplied serially from the topmost to the bottommost check in each field. The bottommost check at each field site received the MD treatment (in order to minimize lateral seepage from adjacent checks), and the upper adjacent check for each field received the FP control treatment. In 2017 and 2019, MD checks were reflooded after 6-7 days of no standing water, while in 2020 MD checks were reflooded once the 0-15 cm surface soil layer reached approximately 35% soil volumetric water content (VWC), which ranged from 6-11 days (Table 3). Across all site-years, the drainage period was initiated an average of 44 days after seeding (DAS) and lasted an average of 7 days. At one site-year (Willows-20), an additional drainage period occurred earlier in the season for herbicide application approximately 24 DAS in which both MD and FP checks were drained and reflooded shortly after.

**Table 3.** Water management practices, including date of drainage and reflood as well as duration of no standing water for each site-year. Drain dates refer to when checks had no standing water. DAS = days after seeding.

| Site-year | Drain Date (DAS) | Reflood Date (DAS) | Duration of no standing water |
| --- | --- | --- | --- |
| Elverta-17 | 7/1 (36) | 7/8 (43) | 7 days |
| Marysville-19 | 7/4 (48) | 7/10 (54) | 6 days |
| Willows-19 | 7/13 (57) | 7/19 (63) | 6 days |
| Arbuckle-20 | 6/6* (34) | 6/13 (41) | 7 days |
| Willows-20 | 6/29** (44) | 7/8 (53) | 9 days |
| Marysville-20 | 6/21 (43) | 6/27 (49) | 6 days |
| Pleasant Grove-20 | 6/25 (45) | 7/6 (56) | 11 days |

\*Water started draining for herbicide application on 6/4 (standing water was present until 6/6)
\*\* Water started draining for herbicide application on 6/26 (standing water was present until 6/29)

Total grain As and Cd, grain yield, and soil moisture were measured in all years of the study, while GHG emissions were measured only in 2017 and 2020. In each year of the study, samples for grain yield and soil moisture were taken at up to four locations throughout each check. Sampling locations for yield and soil moisture were situated approximately 15 m from the edge of each check and roughly equidistant to each other. In the FP control check in 2020, soil moisture was taken at only one of the four locations within each check since variability was expected to be minimal while flooded. Samples for yield were taken at four locations throughout each check in each year of the study. In 2017 and 2020, samples for total grain As and Cd measurement were taken from the same four sampling locations in which samples for yield and soil moisture were taken. In 2019, polyvinyl chloride (PVC) rings, 0.30 m in diameter and 1 m in height, were installed in the MD checks at each site approximately 1 m apart in four separate blocks for grain As and Cd measurements. At Willows-19, 10 MD rings were installed and compared to 8 FP rings, and at Marysville-19, 8 MD rings were installed and compared to 8 FP rings. Continuous flooding was maintained in FP rings at these site-years by manually refilling with water during the drainage period when needed. Example experimental design figures for each year of the study are included in Supplemental Figures 1 and 2.

### 2.4 Soil moisture measurements

Four different soil moisture parameters were measured at the end of the drainage period at each site-year, including gravimetric water content (GWC), soil water potential (SWP), perched water table (PWT), and volumetric water content (VWC). At 0–15 cm depth, VWC was monitored in both years using capacitance sensors (10HS, Decagon Devices Inc., Pullman, WA). Units for VWC were measured as the ratio of water volume to soil volume expressed as a percentage. The sensors were installed vertically in the soil with centers at a soil depth of 7.5 cm, and have a volume of influence of 1 L, which span from 0.5 to 14.5 cm soil depth. Sensors for VWC were installed at four roughly equidistant locations throughout each treatment check within each site, approximately 15 m from the borders of each check. At 0–15 cm depth, SWP was monitored in both years using electrical resistance sensors (Watermark 200SS, Irrometer Co Inc., Riverside, CA). Units for SWP were measured in kilopascals (kPa). The sensors were installed vertically in the soil with the centers at soil depths of 7.5 cm. Sensors for SWP were installed at four roughly equidistant locations throughout each treatment check within each site, approximately 15 m from the borders of each check. In order to measure PWT, a 70 cm long, 5 cm diameter PVC tube perforated with 1 cm diameter holes spaced approximately 2 cm apart was inserted 60 cm deep into the soil in each MD check at four locations after drilling a hole of the exact same diameter. Soil GWC was measured (in the MD treatment and FP control) immediately before the reflooding event of the MD treatment. Soil GWC was determined by taking four samples per treatment check to a depth of 15 cm using a 1.7 cm diameter soil core. Samples were dried at 105 °C until constant weight. Soil GWC (%), was calculated as in Eq. (1):

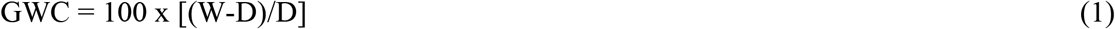

where: W = sample wet weight (g), D = sample dry weight (g).

### 2.5 Greenhouse gas measurements and flux calculations

GHG samples were taken weekly throughout the growing seasons, however, during periods in which emission fluxes were expected to change rapidly (e.g. flooding and drainage events), sampling was conducted in consecutive days or every other day. Gases were captured in cylindrical flux chambers, which consisted of a permanent base installed prior to planting, a variable-height extension to accommodate rice plant growth, and a sealed chamber with a vent tube for pressure equalization (Adviento-Borbe et al., 2013; Pittelkow et al., 2012). Gas samples (25 mL) were taken through silicon septa at 0, 20, 40, and 60 min. and injected into pre-evacuated 12.5-mL glass vials (Labco Ltd., Buckinghamsire, UK). Gas sampling was conducted between 9:00 am and 12:00 pm PST, as soil temperatures and gas fluxes during this time period are expected to be representative of their average daily values (Adviento-Borbe et al., 2013). In order to reduce the effects of intensive gas sampling on rice plants, two collars were installed per block within each check, and sampling alternated between collars for each sampling event. Boardwalks were established prior to planting to minimize soil compaction and to prevent artificially inflated flux values.

All gas samples were analyzed for CH_4_ and N_2_O peak area on a GC-2014 gas chromatograph equipped with a ^63^Ni electron capture detector (ECD) set at 325 °C for N_2_O quantification and a flame ionization detector (FID) for CH_4_ quantification (Shimadzu Scientific, Inst, Columbia, MD, USA). N_2_O was separated by a stainless-steel column packed with Hayesep D, 80/100 mesh at 75 °C. The detection limits of the GC instrument were 1.83 × 10^-4^ mg L^-1^ for both CH_4_ and N_2_O. Results of the GC analyses were accepted if the gas concentrations of CH_4_ and N_2_O standards (1, 3.05, and 9.95 ppm for N_2_O; 1.8, 10.18, and 503 ppm for CH_4_) produced a linear relationship with the voltage output with r^2^ > 0.99. The peak area for each gas sample was then converted to a concentration based on this linear relationship. Fluxes were estimated from the linear increase of gas concentration over time, and gas concentrations were converted to elemental mass per unit area (g ha^-1^ d^-1^) using the Ideal Gas Law with the chamber volume measured at each sampling event, the chamber air temperature measured as each gas sample was taken, and an atmospheric pressure of 0.101 MPa. Fluxes of CH_4_ and N_2_O were computed as:

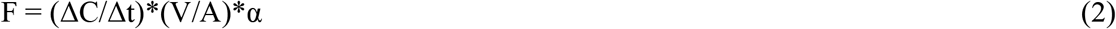

where F is the gas flux rate (g N_2_O–N or CH_4_–C ha^-1^ d^-1^), ΔC/Δt denotes the increase of gas concentration in the chamber (g L^-1^ d^-1^), V is the chamber volume (L), A is area covered by the chamber (ha), and α is a conversion coefficient for elemental C (α = 0.749) or N (α = 0.636). Individual flux values were integrated across all time points via linear interpolation to calculate cumulative seasonal emissions (spring tillage to harvest). As in other similar studies (Adviento-Borbe et al., 2013; Linquist et al., 2015; Pittelkow et al., 2012), gas fluxes with a linear correlation below a predetermined threshold (r^2^ = 0.9) were treated as missing data, and those that were below the GC detection limits were set to zero flux for data analysis.

### 2.6 Grain yield

At physiological maturity, grain yield was measured by manually harvesting four separate 1 m^2^ subplots in each treatment check and control check at each site-year and subsampling approximately 20% of fresh biomass. Grains were oven-dried at 65°C until constant weight, weighed and adjusted to 14% moisture for the estimation of yield. Straw was also oven-dried at 65°C until constant weight and weighed, and harvest index was obtained as the mass ratio of grain to total aboveground biomass.

### 2.7 Grain arsenic and cadmium concentrations

Rice grains were dehulled, polished (white grain samples only), and ball milled to pass a 250 μm sieve. Samples from 2019 and 2020 weighing 0.5 g (two analytical replicates per plot) were digested in 5 mL nitric acid (trace metal grade, 67–70%, VWR International LLC, Radnor, PA, USA) using a microwave digester (Mars 6 Microwave Digestion System, CEM Corporation, Matthews, NC, USA) with a 20 min ramp phase to 180°C and 1600 W at 2450 MHz, which was held for 25 min, followed by a 35 min cool down phase. The 2017 samples were digested in glass digestion tubes by adding 5 mL of nitric acid (trace metal grade, 67–70%, Fisher Chemical, USA) and allowing it to dissolve overnight at room temperature. Samples were further digested in a heating block at 105 °C until the cessation of a brown fog, and then at 120 °C until complete dryness. The ash was re-dissolved with 10 mL of 0.28 M nitric acid. All the digested samples were syringe filtered (0.45 μm), taking care to discard the first 1 mL of the filtrate. The extract was then diluted 5-fold with 18.2 MΩ cm water (Barnstead Nanopure).

Total As and Cd in white and brown grain samples from 2017 were quantified by inductively coupled plasma mass spectrometry (ICP-MS 7900, Agilent Technologies, Santa Clara, CA, USA) with a detection limit of 0.01 μg L^−1^ for both As and Cd. As was monitored at m/ z of 75 and selenium (Se) was also monitored (m/z 77, 78 and 82) to check for polyatomic ^40^Ar^35^Cl interferences on m/z 75. No interferences were observed. Cd was monitored at 112 m/z. Every 10 samples, one blank, one fortified sample, and one certified reference material (1568b, National Institute of Standards and Technology, Gaithersburg, MD, USA) were included as quality control samples. Samples from 2019 and 2020 were also quantified by inductively coupled plasma mass spectrometry (8900 Triple Quadrupole ICP-MS, Agilent Technologies, Santa Clara, CA, USA) with a detection limit of 0.004 μg L^−1^ for As and 0.0018 μg L^−1^ for Cd. Using O_2_ as the reaction cell gas, mass-filtered ^75^As and ^78^Se were measured as reaction product ions ^75^As^16^O^+^ and ^78^Se^16^O^+^, mass-shifted to m/z 91 and 94 respectively, eliminating spectral interferences from doubly charged rare earth elements which overlap As and Se. Cd was monitored at 112 m/ z using He_2_. Every 16 samples, one blank, one fortified sample, and one certified reference material (1568b, National Institute of Standards and Technology, Gaithersburg, MD, USA) were included for quality control. All quality control elements were within the standard quality control criteria, as defined in the FDA Elemental Analysis Manual (FDA, 2012).

### 2.8 Data analysis

GWP was calculated for a 100-yr time horizon using radiative forcing potentials with climate-carbon feedbacks relative to CO_2_ of 28 and 265 for CH_4_ and N_2_O, respectively (Pachauri et al., 2014). We used R studio software (R Core Team, 2022) for analysis of variance on cumulative seasonal CH_4_ emissions, N_2_O emissions, GWP, grain yield, and total grain As and Cd concentrations. Since treatments were not replicated within each site-year, sites were treated as blocks with treatments replicated based on the number of sites in each year. In order to quantify the effect of soil-drying severity on CH_4_ emissions and grain yield, simple linear regression analyses of soil drying parameters, seasonal CH_4_ emissions reductions, and relative grain yield were performed in Excel. The values of soil moisture used for determining the relationship between soil-drying severity and percent reduction in seasonal CH_4_ emissions were measured approximately 3 m from the gas sampling collars at each site-year in which GHG emissions were measured. The values for soil moisture used for determining the relationship between soil-drying severity and relative grain yield were measured in the same locations in which grain yield was harvested for each site-year, approximately 15 m from the edges of each check.

In order to assess check-scale variability in soil-drying during the drainage period (Table 4), locations in which soil moisture parameters were measured alongside a shared levee of a neighboring flooded check were discarded to eliminate the statistical effects of seepage on check-scale drainage. These data points were, however, kept for analysis of the relationship between soil-drying severity and relative grain yield (Figure 1).

**Table 4.** Check-scale mean soil (0-15 cm) moisture parameters [soil water potential (SWP), volumetric water content (VWC), gravimetric water content (GWC), and perched water table (PWT)] measured just before reflood for all site-years and treatments. Numbers in parenthesis represent standard errors of the mean for parameters in which multiple subsamples were taken. In each column of the last two rows, in which mean FP and MD values are presented, means followed by the same letter are not significantly different at P<0.05. In the last two rows, numbers in parenthesis represent standard errors of the mean across all site-years.

| Site-year | Treatment | SWP | VWC | GWC | PWT |
| --- | --- | --- | --- | --- | --- |
|  |  | kPa | % | % | cm |
| Elverta-17 | FP | 0 | 46 | 46 | +13 |
|  | MD | -15 | 45 | 32 (2) | -33 (2) |
| Marysville-19 | FP | 0 | 46 (0) | 48 (1) | +13 |
|  | MD | -20 (6) | 44 (0) | 32 (0) | -31 (4) |
| Willows-19 | FP | 0 | 48 (1) | 47 (1) | +13 |
|  | MD | -25 (2) | 45 (0) | 36 (0) | -24 (9) |
| Arbuckle-20 | FP | 0 | 47 | 48 (0) | +13 |
|  | MD | -46 (17) | 37 (2) | 32 (3) | -29 (14) |
| Willows-20 | FP | 0 | 50 | 52 (2) | +13 |
|  | MD | -70 (6) | 36 (0) | 33 (2) | -19 (0) |
| Marysville-20 | FP | 0 | 45 | 50 (1) | +13 |
|  | MD | -29 (2) | 34 (0) | 25 (2) | -51 (2) |
| Pleasant | FP | 0 | 51 | 58 (1) | +13 |
| Grove-20 | MD | -67 (21) | 34 (2) | 31 (5) | -30 (13) |
| Mean FP |  | 0 (0) a | 48 (1) a | 50 (2) a | +13 (0) a |
| Mean MD |  | -39 (8) b | 39 (2) b | 32 (1) b | -31 (4) b |

**Figure 1.**
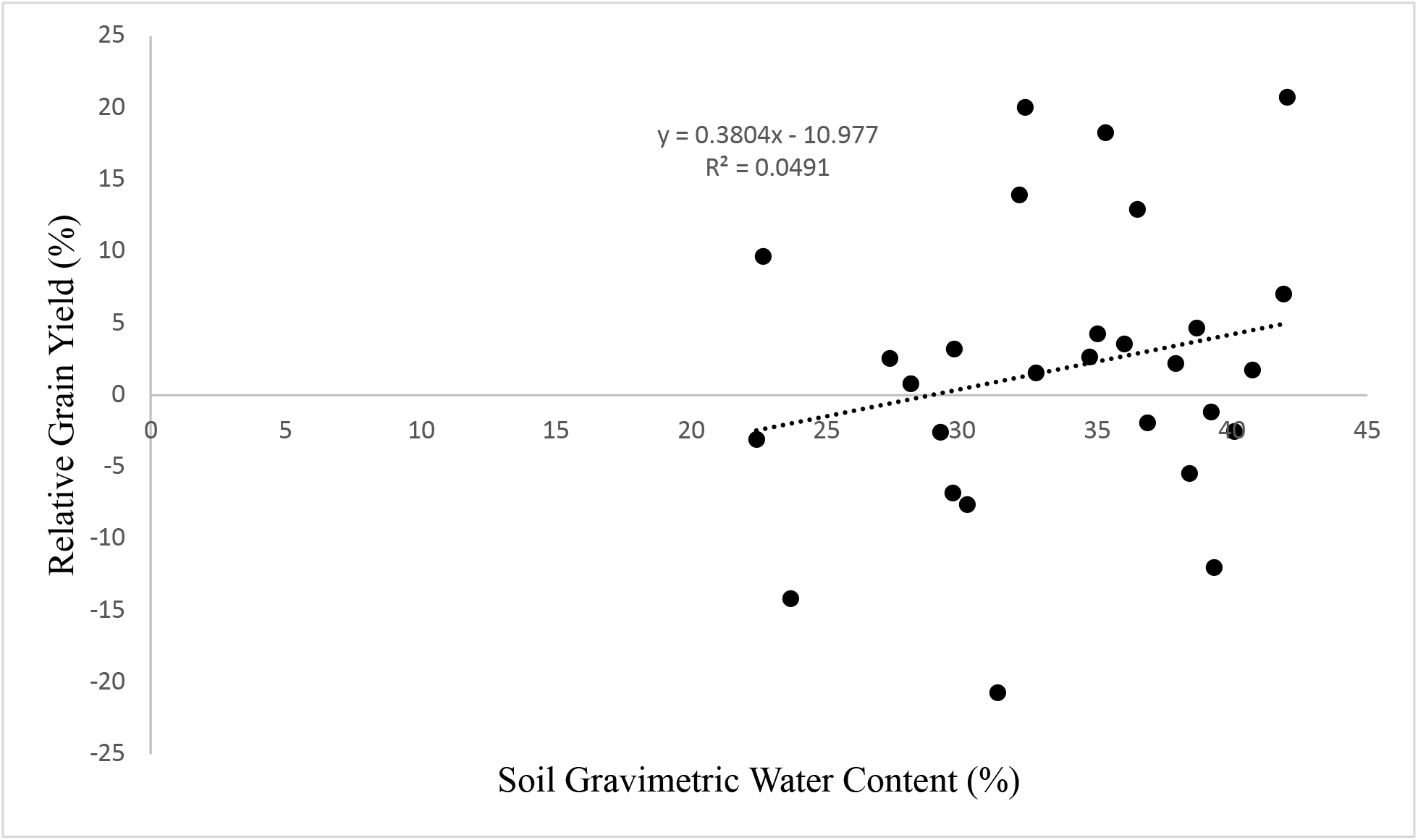
Simple linear regression between soil gravimetric water content (%) measured just before reflood and relative grain yield for all subsamples from drainage treatments compared to the FP control at each site-year.

## 3. Results

### 3.1 Soil moisture

Midseason drainage decreased soil moisture values during the drying period at all site-years in the MD treatments compared to the FP control, as was expected (Table 4). Soil-drying severity, however, was somewhat variable across site-years. Drainage reduced SWP to an average of -39 kPa (range=-15 to -70 kPa), compared to 0 in the FP control across all seven site-years. Drainage reduced VWC to an average of 39% (range=34% to 45%), compared to an average of 48% in the FP control. Additionally, drainage decreased GWC to an average of 32% (range=25% to 36%), compared to an average of 50% in the FP control. The PWT was reduced to an average of 31 cm below the surface (range=19 to 51 cm), compared to an average seasonal floodwater height of 13 cm above the surface for the FP control.

At Arbuckle-20 and Pleasant Grove-20, check-scale soil moisture by the end of the drainage period was somewhat variable. Across site-years in general, check-scale SWP and PWT were most variable while variation in VWC and GWC within each site-year was low (Table 4).

### 3.2 Grain yield

Drainage did not significantly affect grain yield compared to the FP control, however, average grain yield did vary across each site-year (Table 5). Across each site, yields were highest in 2017 and in 2020 (average=12.4 and 12.3 Mg ha^-1^, respectively) and lowest in 2019 (average=11.4 Mg ha^-1^).

**Table 5.** Cumulative seasonal CH_4_ and N_2_O emissions, GWP, and grain yield for each site-year and treatment. Numbers in parentheses represent the standard errors of the mean. In each column of the last two rows, in which mean FP and MD values are presented, means followed by the same letter are not significantly different at P<0.05. In the last two rows, numbers in parenthesis represent standard errors of the mean across all site-years.

| Site-year | Treatment | CH<sub>4</sub><br>kg CH<sub>4</sub>-C<br>ha<sup>-1</sup> | N<sub>2</sub>O<br>kg N<sub>2</sub>O-N<br>ha<sup>-1</sup> | GWP<br>kg CO<sub>2</sub> eq ha<sup>-1</sup> | Yield<br>Mg ha<sup>-1</sup> |
| --- | --- | --- | --- | --- | --- |
| Elverta-17 | FP | 401 (0.1) | -0.02 (0) | 14969 (5) | 12.2 (0.3) |
|  | MD | 186 (15) | 0.12 (0) | 6994 (578) | 12.5 (0.1) |
| Marysville-19 | FP | - | - | - | 10.8 (0.5) |
|  | MD | - | - | - | 11.9 (0.6) |
| Willows-19 | FP | - | - | - | 11.5 (0.1) |
|  | MD | - | - | - | 11.5 (0.1) |
| Arbuckle-20 | FP | 57 (6) | 0 (0) | 2128 (233) | 13.9 (0.1) |
|  | MD | 37 (4) | 0.17 (0.2) | 1458 (93) | 13.7 (0.4) |
| Willows-20 | FP | 136 (56) | 0 (0) | 5070 (2088) | 9.8 (0.4) |
|  | MD | 109 (29) | 0.52 (0.1) | 4302 (1026) | 10.4 (0.9) |
| Marysville-20 | FP | 346 (28) | 0 (0) | 12921 (1036) | 12.2 (0.5) |
|  | MD | 79 (14) | 0.14 (0.1) | 3005 (479) | 12.5 (0.4) |
| Pleasant<br>Grove-20 | FP | 38 (0) | 0 (0) | 1435 (17) | 13.5 (0.1) |
|  | MD | 9 (2) | 0.29 (0.1) | 449 (46) | 12.5 (0.5) |
| Mean FP |  | 196 (87) a | 0 (0) a | 7305 (3267) a | 12.0 (4.5) a |
| Mean MD |  | 84 (38) b | 0.25 (0.1) a | 3242 (1450) b | 12.1 (4.6) a |

Using data from each subsample, a simple linear regression was performed between soil GWC in the MD treatment, measured just before reflood, and grain yield in the MD treatment relative to that of the FP control (Figure 1). This relationship was not significant (P-value = 0.34) and showed that drainage had a minimal effect on grain yield.

### 3.3 Greenhouse gas emissions and global warming potential

Average daily CH_4_ flux of the FP control varied between site-years, the seasonal peak of which ranged between 946 and 9880 g CH_4_-C ha^-1^ day^-1^. The peak for the FP control occurred after the pre-harvest drain (125 DAS) at Elverta-17 (Fig. 2a), while this peak occurred around mid-season for all other site-years, ranging between 18 and 96 DAS. For three site-years (Elverta-17, Willows-20, and Pleasant Grove-20), average daily CH_4_ flux increased immediately after drainage followed by a gradual decrease during the remainder of the drying period. At Marysville-20 and Arbuckle-20, average daily CH_4_ flux decreased immediately after drainage and remained low throughout the drainage period.

**Figure 2a-e.**
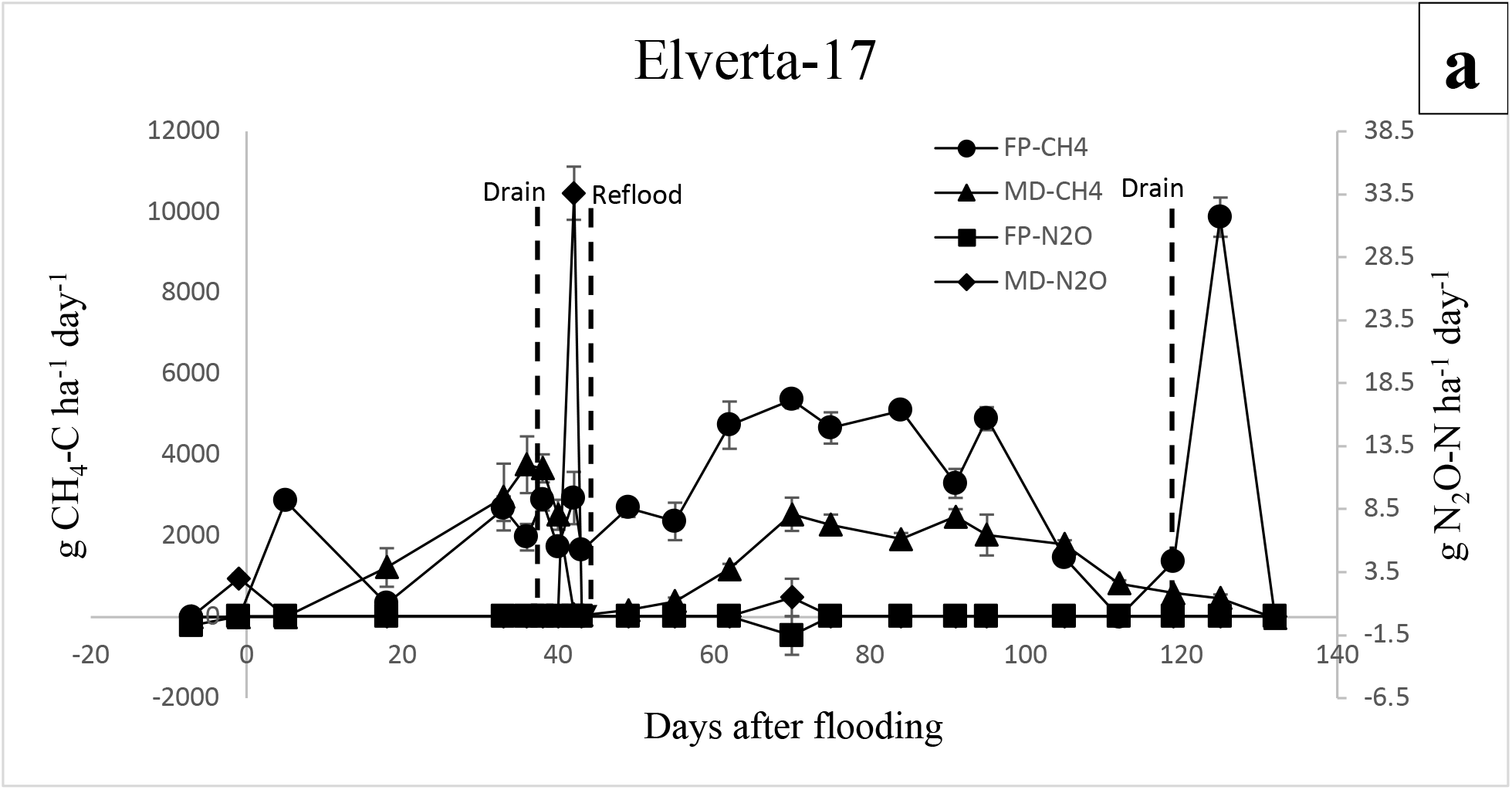
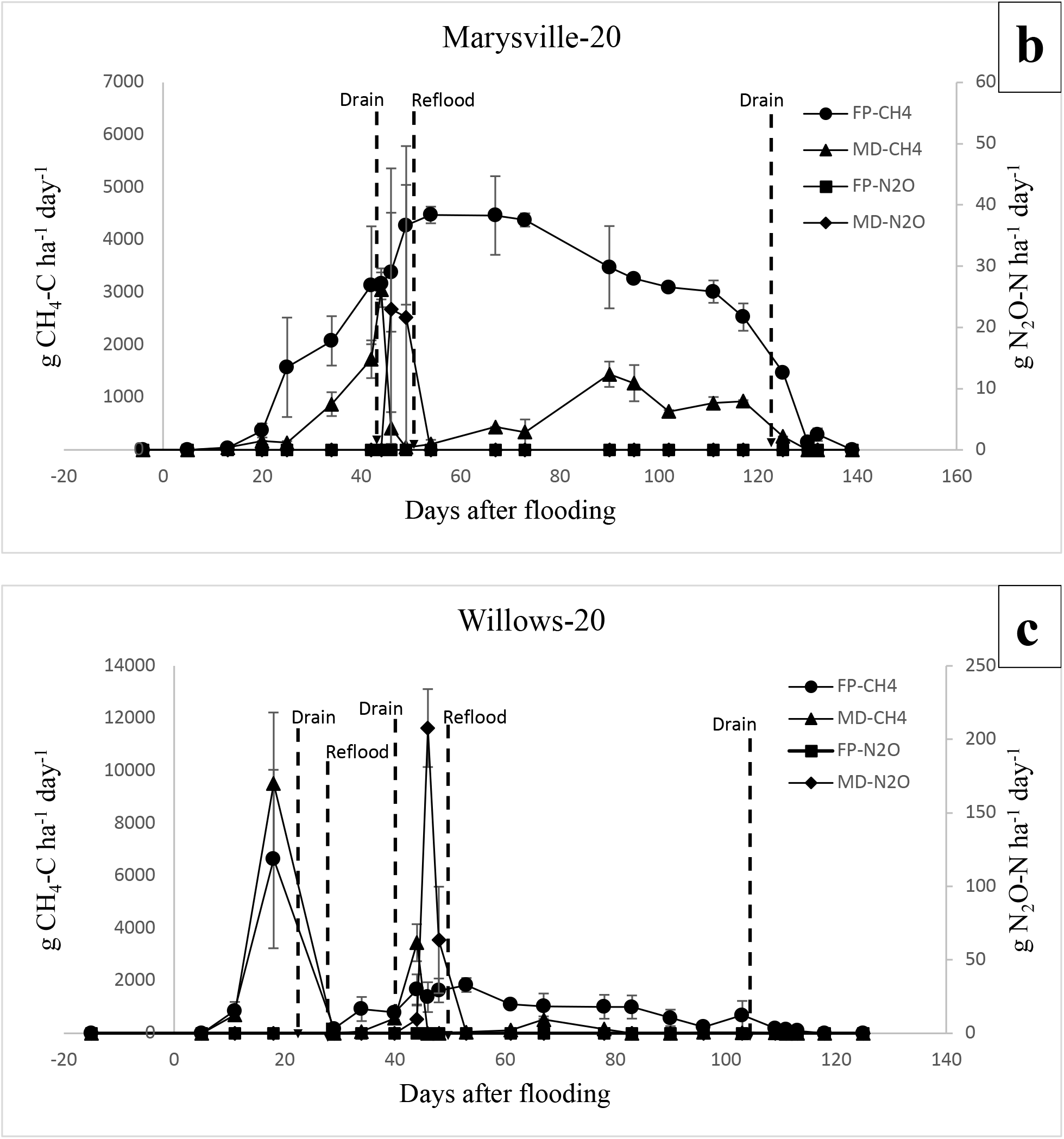
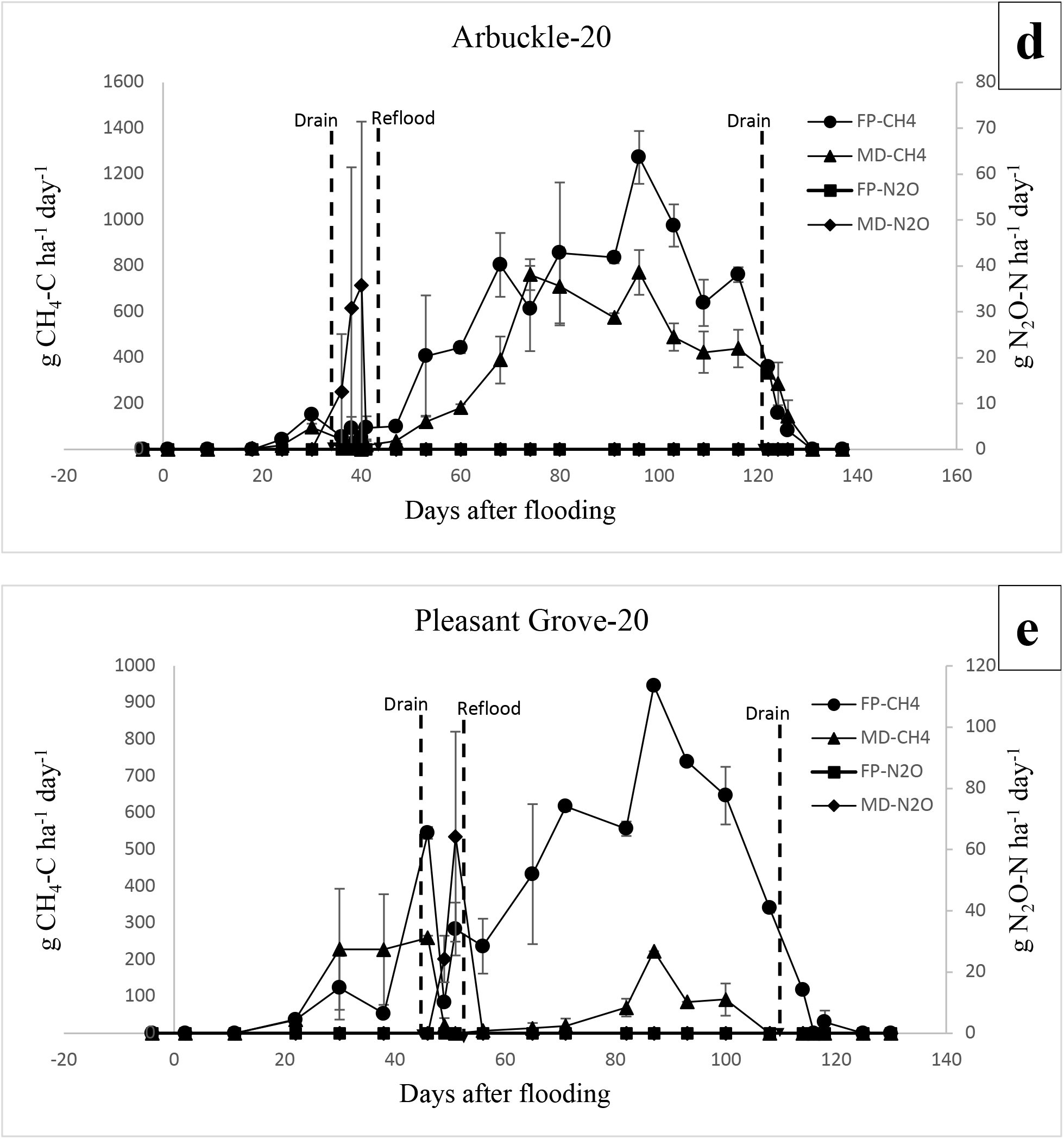
Daily CH_4_ and N_2_O emissions at each site and for each treatment in 2017 and 2020. Dashed vertical lines indicate when drainage periods started and ended for each treatment. Error bars represent the standard errors of the mean.

The imposed drainage period coincided with herbicide application for two site-years. The FP and MD checks at Arbuckle-20 and Willows-20 were both drained for herbicide application 32 and 41 DAS, respectively, which led to a decrease in average daily CH_4_ flux in the FP control at Arbuckle-20 (Fig. 2d). In each instance, FP checks were reflooded after approximately 3 days, while MD checks were allowed to dry for an extended period of time according to the treatment. Additionally, at Willows-20, FP and MD checks were drained for herbicide application 24 DAS, leading to a decrease in average daily CH_4_ flux for both treatments. For all site-years after reflooding, average daily CH_4_ flux of MD checks remained consistently lower than that of FP checks throughout the remainder of the growing season and increased at a slower rate than that of the FP checks. As mentioned earlier at Elverta-17, a high peak in average daily CH_4_ flux from the FP control was observed after the end-of-season drain before harvest (9880 g CH_4_-C ha^-1^ day^-1^). An end-of-season spike was also observed from the FP control at Marysville-20 and Pleasant Grove-20, although these spikes were much smaller (290 and 31 g CH_4_-C ha^-1^ day^-1^, respectively) compared to that of Elverta-17.

Overall, drainage reduced cumulative seasonal CH_4_ emissions by between 20-77% across all five site-years of the study. Locations in which rice straw from the previous growing season was incorporated back into the soil after harvest yielded the highest cumulative seasonal CH_4_ emissions from the FP control (Marysville-20, Elverta-17, and Willows-20) (Table 5). Willows- 20 and Arbuckle-20 experienced the smallest percent reductions in cumulative seasonal CH_4_ emissions compared to the control at 20% and 35%, respectively. This was primarily due to the occurrence of multiple drainage events in the control for herbicide applications at Willows-20 and comparatively low soil-drying severity at Arbuckle-20. Locations in which rice straw was either burned the previous growing season or in which the field was left fallow beforehand yielded the lowest cumulative seasonal CH_4_ emissions in the control (Arbuckle-20 and Pleasant Grove-20) (Table 5).

The results of the simple linear regression between soil GWC measured just before reflood and percent reduction in seasonal CH_4_ emissions compared to the FP control revealed that soil GWC explains approximately 55% of the variation in percent reduction in CH_4_ emissions on-farm (Fig. 3). This relationship also indicates that for every 1% reduction in GWC, seasonal CH_4_ emissions are reduced on-farm by approximately 3.2%, which lies within the 95% confidence interval of the same relationship determined using small on-station plots in Perry et al. (2022) (1.1-3.9%).

**Figure 3.**
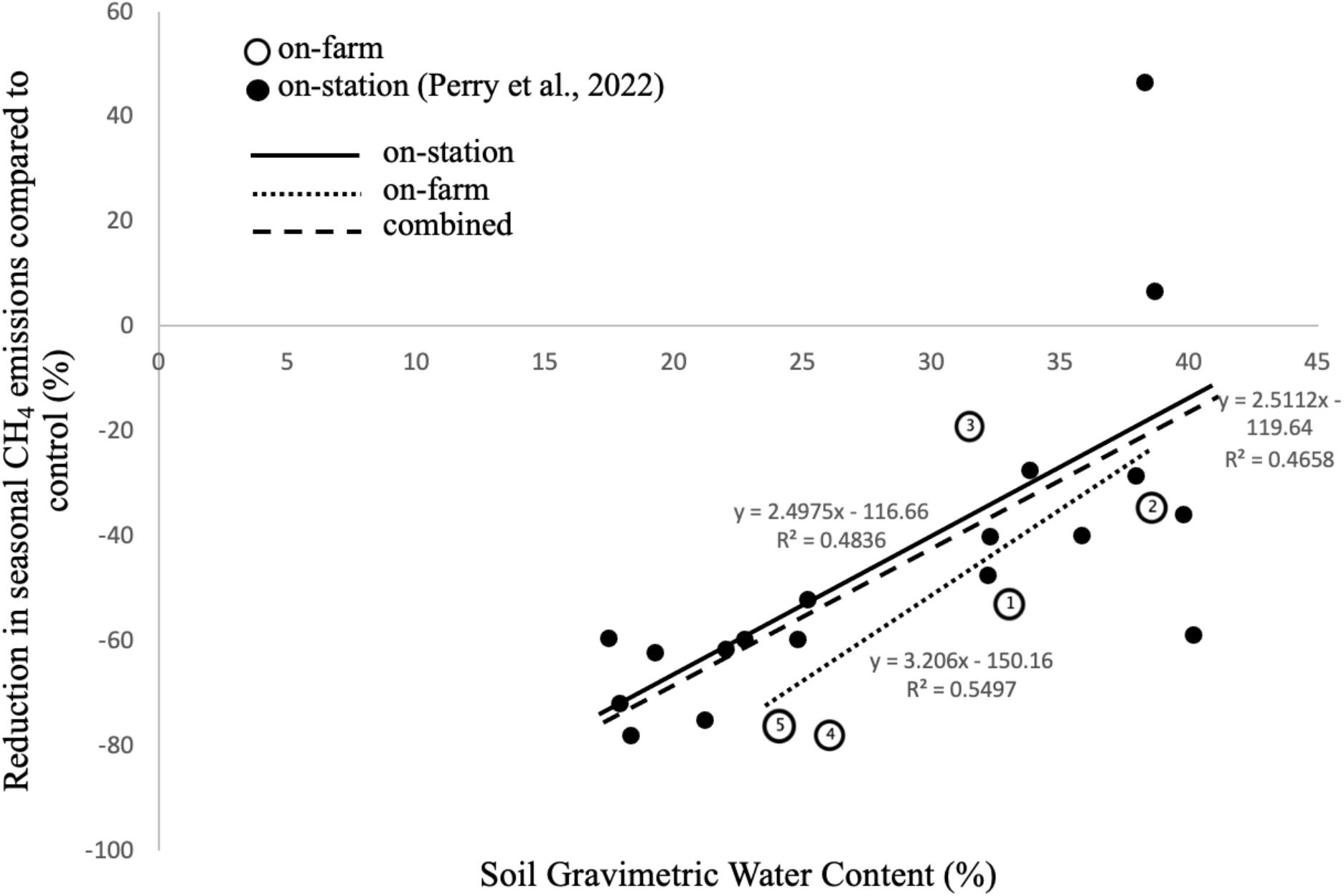
Simple linear regression between soil gravimetric water content (%) measured just before reflood and reduction in seasonal CH_4_ emissions compared to the control (%). Data spans from 2017-2020 and includes single midseason drainage events of varying soil-drying severity and timing tested at the Rice Experiment Station (Perry et al., 2022) as well as single midseason drainage events in conventional rice growers’ fields from this study. Values for seasonal CH_4_ emissions used for the control were the mean values of all control plots in each given year of each study. Open circles represent on-farm data points, and the numbers within each circle correspond to individual site-years as follows: Elverta-17 (1), Arbuckle-20 (2), Willows-20 (3), Marysville-20 (4), Pleasant Grove-20 (5).

With regards to average daily N_2_O flux, spikes in N_2_O emissions occurred primarily during the midseason drainage periods at each site-year (Fig. 2a-e). On average, emissions of N_2_O began between 2-3 days after floodwater subsidence, and average daily N_2_O flux peaked an average of 5 days after drainage, ranging between 22-208 g N_2_O-N ha^-1^ day^-1^ across all site-years. Across all site-years, N_2_O emissions never spiked upon reflooding of the MD treatment. Across all site-years and drainage treatments, cumulative seasonal emissions of N_2_O were low (average=0.248 kg N_2_O-N ha^-1^) and accounted for only 3% of the seasonal global warming potential (GWP) across all drainage treatments. At four of the five site-years in which GHG emissions were measured, cumulative seasonal N_2_O emissions were zero in the FP control, as expected, and at one site-year (Elverta-17), negative seasonal N_2_O emissions were observed in the FP control, suggesting N_2_O uptake.

As yields did not vary considerably between treatments for each site-year of the study and N_2_O emissions were low across treatments, percent reductions in GWP were similar to that of cumulative seasonal CH_4_ emissions. Percent reductions in seasonal GWP compared to the control ranged between 15-77%.

### 3.4 Grain arsenic and cadmium concentrations

Averaged across all site-years, the drainage period significantly decreased total grain As concentration in both white and brown rice (Table 6), while it did not significantly affect grain Cd concentration. At Marysville-20, Willows-20, and Pleasant Grove-20, the greatest decreases in grain As in white rice were seen, with decreases of 19%, 26%, and 47%, respectively, compared to the FP control. On average, drainage decreased total grain As by approximately 23% in white rice across all site-years as well as approximately 20%, on average, in brown rice. Surprisingly, drainage also decreased average grain Cd concentration in both white and brown rice across all site years, although not significantly so.

**Table 6.** Grain arsenic and cadmium concentrations from polished white rice and brown rice for each site-year and treatment. Numbers in parenthesis represent standard errors of the mean. In each column of the last two rows, in which mean FP and MD values are presented, means followed by the same letter are not significantly different at P<0.05.

| Site | Treatment | White | Brown |
| --- | --- | --- | --- |

|  |  | As | Cd | As | Cd |
| --- | --- | --- | --- | --- | --- |
|  |  | ug/kg grain |  |  |  |
| Elverta-17 | FP | 0.139 (0.01) | 0.005 (0) | 0.293 (0.02) | 0.006 (0) |
|  | MD | 0.119 (0.01) | 0.006 (0) | 0.251 (0.01) | 0.006 (0) |
| Marysville-19 | FP | n/a | n/a | 0.239 (0.01) | 0.006 (0) |
|  | MD | n/a | n/a | 0.195 (0.03) | 0.008 (0) |
| Willows-19 | FP | n/a | n/a | 0.145 (0) | 0.005 (0) |
|  | MD | n/a | n/a | 0.136 (0) | 0.006 (0) |
| Arbuckle-20 | FP | 0.116 (0) | 0.019 (0) | 0.145 (0) | 0.019 (0) |
|  | MD | 0.115 (0) | 0.014 (0) | 0.142 (0.01) | 0.014 (0) |
| Willows-20 | FP | 0.151 (0.01) | 0.005 (0) | 0.203 (0.02) | 0.005 (0) |
|  | MD | 0.112 (0.01) | 0.004 (0) | 0.155 (0.02) | 0.004 (0) |
| Marysville-20 | FP | 0.166 (0) | 0.005 (0) | 0.206 (0.01) | 0.006 (0) |
|  | MD | 0.134 (0) | 0.006 (0) | 0.177 (0) | 0.006 (0) |
| Pleasant | FP | 0.178 (0.02) | 0.007 (0) | 0.235 (0.02) | 0.007 (0) |
| Grove-20 | MD | 0.094 (0.01) | 0.009 (0) | 0.119 (0.01) | 0.008 (0) |
| Mean FP |  | 0.150 (0.01) a | 0.820e-2 (0) a | 0.206 (0.01) a | 0.734e-2 (0) a |
| Mean MD |  | 0.115 (0) b | 0.777e-2 (0) a | 0.166 (0.01) b | 0.720e-2 (0) a |

## 4. Discussion

### 4.1 Grain yield

Across all site-years of this study, grain yield was not significantly different in the MD treatment compared to the FP control (Table 5). Similar findings have been illustrated in previous studies in which drainage periods can maintain or even increase rice grain yield as a result of increased root oxidation and cytokinin activity (Yang et al., 2017) or increased sugar-to-starch conversion in grains and photosynthetic rates (Zhang et al., 2009). In this study, a mechanistic explanation behind the lack of reduction in grain yields under drainage conditions compared to continuous flooding across all site-years is difficult to ascertain. However, it is likely that soil-drying severity and the duration of the drainage periods were not great enough to cause significant impacts on yields.

Yields, however, did vary somewhat across site-years (range = 9.8-13.9 Mg ha^-1^), with the lowest yields seen at Willows-20 and the highest at Arbuckle-20. Relatively low yields in the MD treatment and the FP control at Willows-20 (average=10.1 Mg ha^-1^) compared to the other sites can be explained by moderate weed infestation in this field during this year, whereas proliferation of weeds was not a factor at any of the other site-years.

### 4.2 Methane emissions

Cumulative seasonal CH_4_ emissions varied between locations and years in this study. Variation in cumulative seasonal CH_4_ emissions from the FP control across locations can be explained primarily by the different straw management practices employed at each location in the previous growing season as well as differences in soil organic carbon and soil texture. In flooded rice systems, early peaks in seasonal CH_4_ emissions are often attributed to decomposition of the previous crop residues, as residues left in the field from the previous growing season can provide additional carbon substrate for methanogenesis (Chidthaisong and Watanabe, 1997; Yan et al., 2005). At three site-years in this study (Elverta-17, Marysville-20, and Willows-20), rice straw from the previous growing season was chopped and incorporated into the soil, followed by winter flooding, and at these three site-years, cumulative seasonal CH_4_ emissions from the FP control were highest (range=136-401 kg CH_4_-C ha^-1^) across all site-years. Alternatively, site-years in which the field location was fallowed in the previous growing season (Pleasant Grove-20) or in which rice straw was burned before the season (Arbuckle-20) resulted in the lowest cumulative seasonal CH_4_ emissions from the FP control (38 and 57 kg CH_4_-C ha^-1^, respectively).

Additionally, in a meta-analysis conducted by Linquist et al. (2018), it was found that high clay content generally decreases cumulative seasonal CH_4_ emissions from flooded rice systems. This is consistent with results from this study in that the sites with the highest soil clay content (Pleasant Grove-20 and Arbuckle-20) had the lowest cumulative seasonal CH_4_ emissions of all site-years in the FP control (38 and 57 kg CH_4_-C ha^-1^, respectively).

With regards to the effectiveness of the drainage period for GHG mitigation, seasonal CH_4_ emissions were further decreased with increasing soil-drying severity during the drainage period (Fig. 3). Data combined from Perry et al. (2022), in which a similar single midseason drain was employed in rice systems, showed that soil GWC measured just before reflooding explains a significant portion of the variation in percent reduction of CH_4_ emissions. The site-years of this study in which a midseason drainage period was also employed showed a similarly strong relationship between soil-drying and reduction in CH_4_ emissions as that of plots studied in Perry et al. (2022). This relationship also indicates that maximum reductions in CH_4_ emissions may likely occur between 20-25% GWC, and that further drying past this point may not achieve further reductions in CH_4_ emissions but instead may pose a risk for yield reduction.

Additionally, understanding the relationship between soil-drying and CH_4_ emissions reductions may be useful in predicting seasonal CH_4_ emissions reductions from rice fields in which a midseason drain is implemented. Drying fields for an average duration of 7 days and reaching approximately 32% soil GWC by the end of the drainage period may reduce cumulative seasonal CH_4_ emissions by approximately 40% according to Figure 3. Establishing this relationship could be a useful indicator of the magnitude of CH_4_ emissions reductions for rice fields in which midseason drainage is practiced. Importantly, and as reported with regards to variation in soil-drying severity, soil GWC was not highly variable throughout each MD check (Table 4), and therefore it is possible that CH_4_ emissions reductions from a single check and even whole fields may be relatively uniform if surface floodwater is properly and effectively removed from a field during the drainage period.

Reductions in seasonal CH_4_ emissions at the site-years of this study were approximately 20%, 35%, 54%, 77%, and 76% (average = 52%) with the 20% reduction coming from a site-year in which the MD check and FP control plot were drained twice for herbicide application (Willows-20), thereby reducing the effect of drainage compared to the control. However, on average, the reduction in seasonal CH_4_ emissions observed in this study was higher than those previously reported for single drainage events, both in a U.S.-based meta-analysis by Linquist et al. (2018) (average = 39%; 95% confidence interval 30-47%) and a global meta-analysis by Jiang et al. (2019) (average = 33%; 95% confidence interval 12-49%). Although soil-drying severity was somewhat variable across site-years in this study, the large percent reductions in seasonal CH_4_ emissions observed can be attributed to the relatively high soil-drying severity, on average, of the drainage periods in this study compared to other studies.

### 4.3 Nitrous oxide emissions and GWP

Overall, cumulative seasonal N_2_O emissions were low (average = 0.248 kg N_2_O-N ha^-1^) across all locations and drainage treatments. Emissions of N_2_O primarily occurred during the drainage periods for MD treatments and accounted for approximately 3% of the total GWP across all drainage treatments. Cumulative seasonal N_2_O emissions were highest in the MD treatment at Willows-20 (0.52 kg N_2_O-N ha^-1^), which can likely be explained by the relatively high rate of mineral N fertilizer applied at this site-year (210 kg ha^-1^) and total soil N content (0.18%) at this site compared to the other site-years of this study.

At one site-year (Elverta-17), negative seasonal N_2_O emissions were observed, which suggests the occurrence of N_2_O uptake and can be somewhat common in flooded rice fields (Majumdar, 2013). Although the mechanisms of N_2_O uptake in flooded rice soils are not widely studied and factors affecting N_2_O uptake can be quite variable, Liu et al. (2007) reported that rice straw from the previous growing season can increase the magnitude of negative N_2_O fluxes. This may explain the negative N_2_O fluxes observed at Elverta-17, which was one of the three site-years in which rice straw was chopped and incorporated into the soil prior to the growing season.

As yields did not vary significantly between treatments for each site-year of this study and N_2_O emissions were low across treatments, reductions in GWP were similar to that of cumulative seasonal CH_4_ emissions, with percent reductions in seasonal GWP ranging between 15-77%, compared to the FP control.

### 4.4 Grain arsenic and cadmium concentrations

In general, at site-years in which soil-drying severity was highest during the drainage period, the largest percent decreases in total grain As concentration were observed, which is consistent with previous findings (Carrijo et al., 2019). On average, drainage significantly decreased total grain As in white and brown rice by approximately 20%, compared to the FP control (Table 6). This reduction is, however, lower than that of previous studies in which total grain As concentrations were reduced by between 41-68% in this region across a range of soil-drying severities and number of drainage events (Carrijo et al., 2018; Carrijo et al., 2019).

Non-continuous flooding practices such as AWD have been shown to decrease grain As but increase Cd concentrations in rice grains (Makino et al., 2016; Norton et al., 2017; Yang et al., 2017). Flooding-induced soil anaerobic conditions favor As mobilization due to the reduction of As(V) to more labile As(III) and the reductive dissolution of As-bearing Fe(III) solid phases (Fendorf et al., 2007). By contrast, anaerobic conditions can limit grain Cd because microbially mediated sulfate reduction can form poorly soluble Cd sulfide, and flooding can neutralize acidic soils, promoting Cd^2+^ adsorption to soil solids (Wang et al., 2019). When drainage occurs, these Cd immobilization processes are reversed, enhancing Cd bioavailability for uptake by rice (Rinklebe et al., 2016). Hence, it was expected that drainage would significantly increase grain Cd concentration, averaged across all site-years, compared to the FP control, however, this was found not to be the case. Although drainage increased grain Cd concentration for 3 out of 5 site-years with white rice (Elverta-17, Marysville-20, and Pleasant Grove-20) and for 3 out of 7 site-years with brown rice (Marysville-19, Willows-19, and Pleasant Grove-20), drainage actually decreased grain Cd in white and brown rice on average across all site-years, although this effect was not significant. One explanation for limited Cd uptake may be residual levels of soil ammonium still present at the time of drainage, which has been shown to inhibit Cd uptake in rice (Wu et al., 2018). Additionally, a single midseason drainage event, rather than multiple drainage events, may not have been sufficient to increase grain Cd concentration, even if Cd was temporarily mobilized in the soil solution during drainage.

## 5. Conclusion

In this study, it was found that the implementation of a single midseason drainage event has the potential to decrease seasonal cumulative CH_4_ emissions on-farm from flooded rice cultivation while maintaining peak agronomic production in California rice systems. Soil-drying severity during the drainage period was strongly related to reductions in CH_4_ emissions, and drainage had no significant impact on grain yield. Emissions of N_2_O due to drainage were low and were not a significant component of seasonal GWP. Additionally, drainage significantly decreased total grain As concentration in both white and brown rice without significantly affecting grain Cd concentration, compared to the control. Peak seasonal CH_4_ emissions varied across site-years representing different soil types and straw management practices, which further illustrates the importance of the continued assessment of the effectiveness of midseason drainage across a wide range of soil types, farmer management practices, and environmental conditions. There are several obstacles to widespread adoption of midseason drainage and non-continuous flooding practices as a whole, particularly in California, however, this research highlights the potential of midseason drainage to bring about significant environmental and human health benefits.

## References

Adviento-Borbe, Maria Arlene, et al. “Optimal fertilizer nitrogen rates and yield-scaled global warming potential in drill seeded rice.” Journal of environmental quality 42.6 (2013): 1623–1634.

Balaine, Nimlesh, et al. “Greenhouse Gases from Irrigated Rice Systems under Varying Severity of Alternate-Wetting and Drying Irrigation.” Soil Science Society of America Journal83.5 (2019): 1533–1541.

Barker, Randolph; Molle F. Evolution of irrigation in South and Southeast Asia. Research Report 5. 2004. pp. 1–46. Available: http://www.iwmi.cgiar.org/assessment/files/pdf/publications/ResearchReports/CARR5.pdf

Bateman, E. J., and E. M. Baggs. “Contributions of nitrification and denitrification to N 2 O emissions from soils at different water-filled pore space.” Biology and fertility of soils 41.6 (2005): 379–388.

Bhowmick, Subhamoy, et al. “Arsenic in groundwater of West Bengal, India: a review of human health risks and assessment of possible intervention options.” Science of the Total Environment 612 (2018): 148–169.

Carrijo, Daniela R., Mark E. Lundy, and Bruce A. Linquist. “Rice yields and water use under alternate wetting and drying irrigation: A meta-analysis.” Field Crops Research 203 (2017): 173–180.

Carrijo, Daniela R., et al. “Impacts of variable soil drying in alternate wetting and drying rice systems on yields, grain arsenic concentration and soil moisture dynamics.” Field Crops Research 222 (2018): 101–110.

Carrijo, Daniela R., et al. “Irrigation management for arsenic mitigation in rice grain: Timing and severity of a single soil drying.” Science of the total environment 649 (2019): 300–307.

Chidthaisong, Amnat, and Iwao Watanabe. “Methane formation and emission from flooded rice soil incorporated with 13C-labeled rice straw.” Soil Biology and Biochemistry 29.8 (1997): 1173–1181.

Childs, Nathan, Sharon Skorbiansky Raszap, and William D. McBride. “US Rice Production Changed Significantly in the New Millennium, but Remained Profitable.” Amber Waves: The Economics of Food, Farming, Natural Resources, and Rural America 2020.1490-2020–1031 (2020).

CIMIS (California Irrigation Management Information System), 2021. Station Reports. (Accessed on 5 July, 2021 at https://cimis.water.ca.gov/WSNReportCriteria.aspx)

Conrad, R., 2007. Microbial ecology of methanogens and methanotrophs. Adv. Agron. 96, 1–63.

de Vries, Michiel E., et al. “Rice production with less irrigation water is possible in a Sahelian environment.” Field Crops Research 116.1-2 (2010): 154–164.

Jessica Dittmar, Andreas Voegelin, Linda C. Roberts, Stephan J. Hug, Ganesh C. Saha, M. Ashraf Ali, A. Borhan M. Badruzzaman, and Ruben Kretzschmar. “Spatial Distribution and Temporal Variability of Arsenic in Irrigated Rice Fields in Bangladesh”. Environmental Science & Technology, 2007 41 (17): 5967–5972. DOI: 10.1021/es0702972.

Dunn, B. W., and D. S. Gaydon. “Rice growth, yield and water productivity responses to irrigation scheduling prior to the delayed application of continuous flooding in south-east Australia.” Agricultural Water Management 98.12 (2011): 1799–1807.

FDA, US. “Arsenic Speciation in Rice and Rice Products Using High Performance Liquid ChromatographyInductively Coupled Plasma-Mass Spectrometric Determination.” (2012).

Fendorf, S., et al. “Biogeochemical processes controlling the cycling of arsenic in soils and sediments.” Biophysico-chemical processes of heavy metals and metalloids in soil environments (2007): 313–338.

Hill, J. E., et al. “Rice irrigation systems for tailwater management.” Leaflet-University of California, Cooperative Extension Service (USA) (1991).

IPCC, 2013. Climate change 2013: the physical science basis. In T. F. Stocker et al. (eds.). Working Group I Contribution to the Fifth Assessment Report of the Intergovernmental Panel on Climate Change (pp. 1535). Cambridge, UK, and New York, NY: Cambridge University Press.

Jiang, Yu, et al. “Water management to mitigate the global warming potential of rice systems: A global meta-analysis.” Field Crops Research 234 (2019): 47–54.

Khan, Shahbaz, et al. “Can irrigation be sustainable?.” Agricultural Water Management 80.1-3 (2006): 87–99.

LaHue, Gabriel T., et al. “Alternate wetting and drying in high yielding direct-seeded rice systems accomplishes multiple environmental and agronomic objectives.” Agriculture, ecosystems & environment 229 (2016): 30–39.

Li, Chongyang, et al. “Impact of alternate wetting and drying irrigation on arsenic uptake and speciation in flooded rice systems.” Agriculture, Ecosystems & Environment 272 (2019): 188–198.

Linquist, B.A., K.J. van Groenigen, M.A. Adviento-Borbe, C. Pittelkow, and C. van Kessel. (2012). An agronomic assessment of greenhouse gas emissions from major cereal crops. Glob. Change Biol. 18:194–209.

Linquist, Bruce A., et al. “Reducing greenhouse gas emissions, water use, and grain arsenic levels in rice systems.” Global change biology 21.1 (2015): 407–417.

Linquist, Bruce A., et al. “Greenhouse gas emissions and management practices that affect emissions in US rice systems.” Journal of environmental quality 47.3 (2018): 395–409.

Liu, Xiaojing, et al. “Characteristics of CO 2, CH 4 and N 2 O emissions from winter-fallowed paddy fields in hilly areas of South China.” Frontiers of Agriculture in China 1.4 (2007): 418–423.

Lu, Jun, Taiichiro Ookawa, and Tadashi Hirasawa. “The effects of irrigation regimes on the water use, dry matter production and physiological responses of paddy rice.” Plant and soil223.1-2 (2000): 209–218.

Majumdar, Deepanjan. “Biogeochemistry of N2O uptake and consumption in submerged soils and rice fields and implications in climate change.” Critical reviews in environmental science and technology 43.24 (2013): 2653–2684.

Makino, Tomoyuki, et al. “Simultaneous decrease of arsenic and cadmium in rice (Oryza sativa L.) plants cultivated under submerged field conditions by the application of iron-bearing materials.” Soil Science and Plant Nutrition 62.4 (2016): 340–348.

Martínez-Eixarch, Maite, et al. “Multiple environmental benefits of alternate wetting and drying irrigation system with limited yield impact on European rice cultivation: The Ebre Delta case.” Agricultural Water Management 258 (2021): 107164.

Meharg, Andrew A. “Arsenic in rice–understanding a new disaster for South-East Asia.” Trends in plant science 9.9 (2004): 415–417.

Nalley, Lanier, et al. The Economic Viability of Alternative Wet Dry (AWD) Irrigation in Rice Production in the Mid-South. No. 1374-2016-109340. 2014.

Norton, Gareth J., et al. “Impact of alternate wetting and drying on rice physiology, grain production, and grain quality.” Field Crops Research 205 (2017): 1–13.

Pachauri, Rajendra K., et al. Climate change 2014: synthesis report. Contribution of Working Groups I, II and III to the fifth assessment report of the Intergovernmental Panel on Climate Change. IPCC, 2014.

Pandey, Arjun, et al. “Organic matter and water management strategies to reduce methane and nitrous oxide emissions from rice paddies in Vietnam.” Agriculture, ecosystems & environment 196 (2014): 137–146.

Perry, Henry, Daniela Carrijo, and Bruce Linquist. “Single midseason drainage events decrease global warming potential without sacrificing grain yield in flooded rice systems.” Field Crops Research 276 (2022): 108312.

Pittelkow, C. M., et al. “Agronomic productivity and nitrogen requirements of alternative tillage and crop establishment systems for improved weed control in direct-seeded rice.” Field Crops Research 130 (2012): 128–137.

R Core Team (2022). R: A language and environment for statistical computing. R Foundation for Statistical Computing, Vienna, Austria. URL http://www.R-project.org/.

Rinklebe, Jörg, Sabry M. Shaheen, and Kewei Yu. “Release of As, Ba, Cd, Cu, Pb, and Sr under pre-definite redox conditions in different rice paddy soils originating from the USA and Asia.” Geoderma 270 (2016): 21–32.

Sander, Bjoern Ole, et al. “Climate-based suitability assessment for alternate wetting and drying water management in the Philippines: a novel approach for mapping methane mitigation potential in rice production.” Carbon management 8.4 (2017): 331–342.

Towprayoon, S., K. Smakgahn, and S. Poonkaew. “Mitigation of methane and nitrous oxide emissions from drained irrigated rice fields.” Chemosphere 59.11 (2005): 1547–1556.

UCANR. About California Rice. University of California Agronomy Information and Research Center. (Accessed 09.03.21). http://rice.ucanr.edu/About_California_Rice/

USDA, 2015. Southeast Asia: 2015/16 Rice Production Outlook at Record Levels. USDA Foreign Agricultural Service Commodity Intelligence Report. (Accessed 08.27.21). https://ipad.fas.usda.gov/highlights/2015/06/Southeast_Asia/Index.htm

Wang, Jing, et al. “Iron–manganese (oxyhydro) oxides, rather than oxidation of sulfides, determine mobilization of Cd during soil drainage in paddy soil systems.” Environmental science & technology 53.5 (2019): 2500–2508.

Wu, Zhichao, et al. “Increasing ammonium nutrition as a strategy for inhibition of cadmium uptake and xylem transport in rice (Oryza sativa L.) exposed to cadmium stress.” Environmental and experimental botany 155 (2018): 734–741.

Yan, Xiaoyuan, et al. “Statistical analysis of the major variables controlling methane emission from rice fields.” Global Change Biology 11.7 (2005): 1131–1141.

Yang, Jianchang, Qun Zhou, and Jianhua Zhang. “Moderate wetting and drying increases rice yield and reduces water use, grain arsenic level, and methane emission.” The Crop Journal5.2 (2017): 151–158.

Yao, Fengxian, et al. “Agronomic performance of high-yielding rice variety grown under alternate wetting and drying irrigation.” Field crops research 126 (2012): 16–22.

Zhang, Hao, et al. “An alternate wetting and moderate soil drying regime improves root and shoot growth in rice.” Crop Science 49.6 (2009): 2246–2260.

Zou, Jianwen, et al. “Quantifying direct N2O emissions in paddy fields during rice growing season in mainland China: dependence on water regime.” Atmospheric Environment41.37 (2007): 8030–8042.

