## Supplemental Figures 1 and 2 for "On-farm Implementation of Midseason Drainage to Decrease Greenhouse Gas Emissions and Grain Arsenic Concentration in Rice Systems"

Supplemental Figure 1. Example experimental design for 2019 sites. Samples for grain yield and soil moisture (including GWC, VWC, and SWP) were taken at each numbered FP location in each FP control check. For each MD check, samples for grain yield and soil moisture (including GWC, VWC, SWP, and PWT) were taken at each numbered MD location. Numbered FP and MD locations were situated approximately 15 m from the edge of each check and roughly equidistant to each other. Open circles on the diagram in the MD check represent locations in which samples for grain arsenic and cadmium concentrations were taken in 2019. These samples were taken from rings, 0.30 m in diameter and 1 m in height, placed directly in the soil during the growing season and were situated approximately 1 m apart from each other in four separate blocks with one FP control ring and one MD treatment ring within each block.

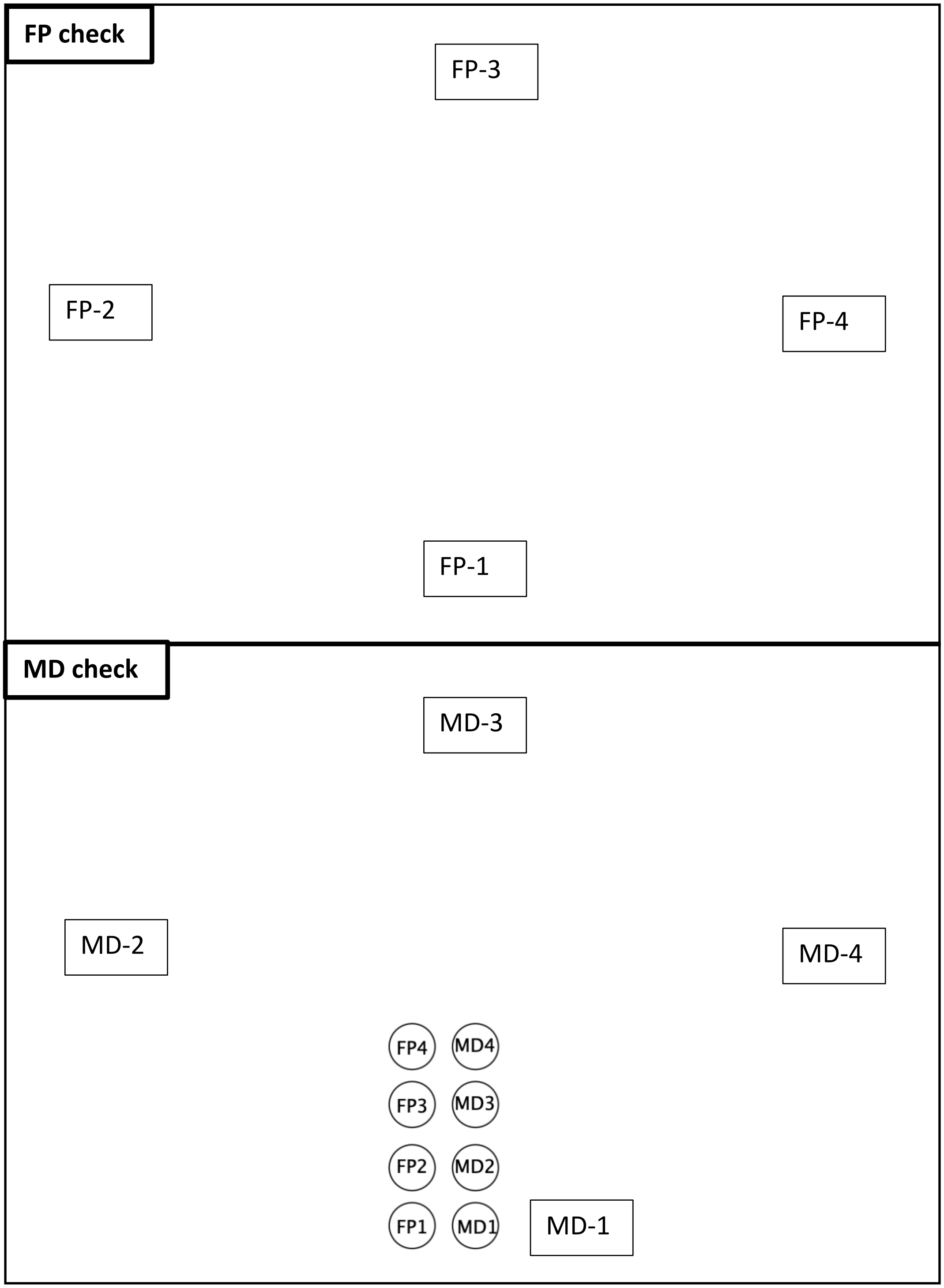

Supplemental Figure 2. Example experimental design for 2017 and 2020 site-years. Samples for grain yield, arsenic, and cadmium concentrations were taken at each numbered FP location in each FP control check. Within each FP check, samples for soil moisture (including GWC, VWC, and SWP) were measured at FP-1 only. For each MD check, samples for grain yield, soil moisture (including GWC, VWC, SWP, and PWT), and grain arsenic and cadmium concentrations were taken at each numbered MD location. Numbered FP and MD locations were situated approximately 15 m from the edge of each check and roughly equidistant to each other. Open circles on the diagram represent locations in which GHG emissions were sampled from collars which were placed directly into the soil prior to planting. During each gas sampling event, GHG samples were taken from two subsamples (i.e. A1 and A2) within each check, and sampling alternated between collars A and B for each gas sampling event to minimize the effects of intensive sampling.

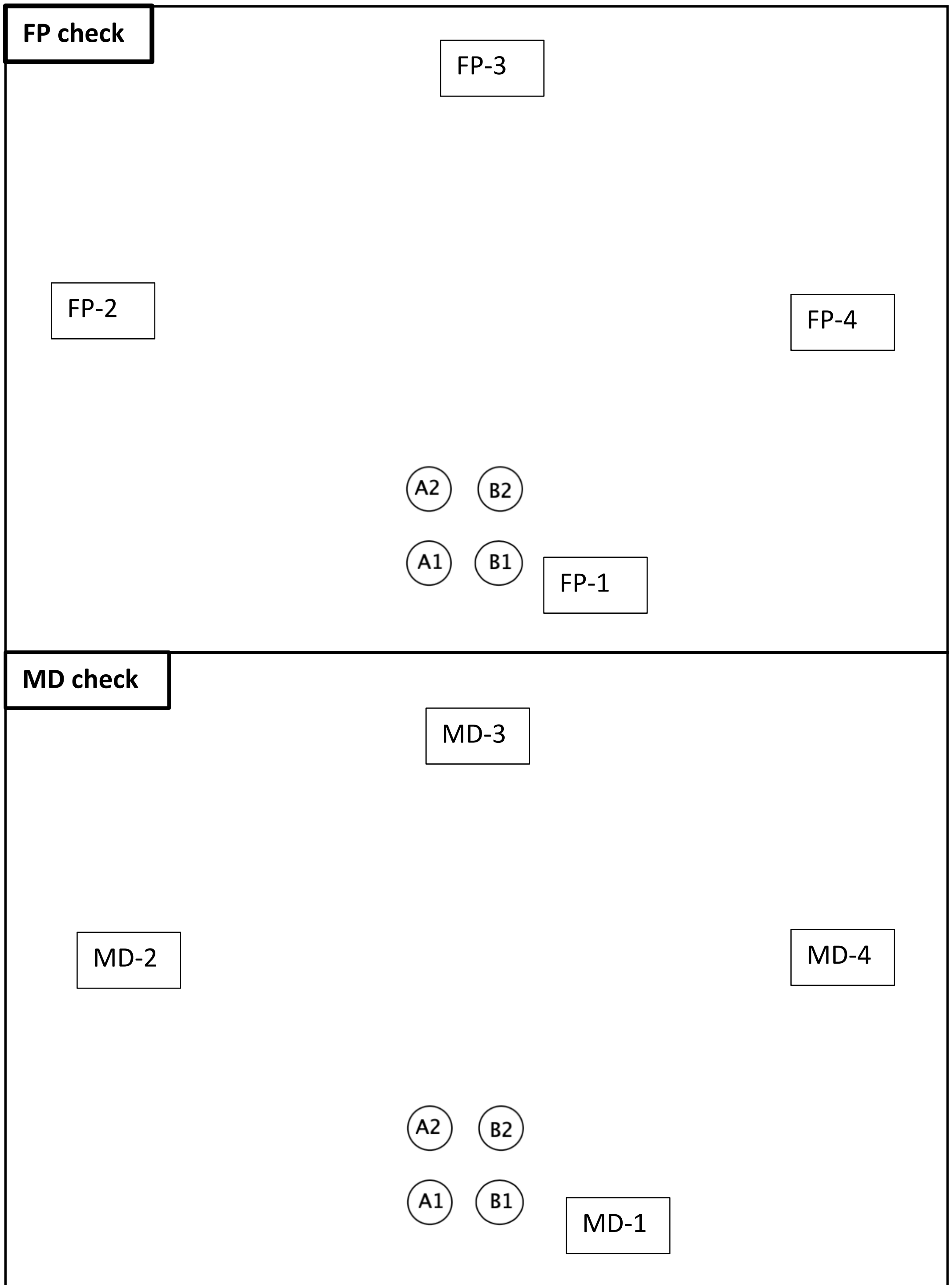
